# Characterization of a rationally engineered *phaCAB* operon with a hybrid promoter design

**DOI:** 10.1101/006106

**Authors:** Iain Bower, Bobby Wenqiang Chi, Matthew Ho Wai Chin, Sisi Fan, Margarita Kopniczky, Jemma Pilcher, James Strutt, Richard Kelwick, Alexander Webb, Kirsten Jensen, Guy-Bart Stan, Richard Kitney, Paul Freemont

## Abstract

Biopolymers, such as poly-3-hydroxy-butyrate (P(3HB)) are produced as a carbon store in an array of organisms and exhibit characteristics which are similar to oil-derived plastics, yet have the added advantages of biodegradability and biocompatibility. Despite these advantages, P(3HB) production is currently more expensive than the production of oil-derived plastics, and therefore more efficient P(3HB) production processes are required. In this study, we describe the model-guided design and experimental characterization of several engineered P(3HB) producing operons. In particular, we describe the characterization of a novel hybrid *phaCAB* operon that consists of a dual promoter (native and J23104) and RBS (native and B0034) design. P(3HB) production was around six-fold higher in hybrid *phaCAB* engineered *Escherichia coli* in comparison to *E. coli* engineered with the native *phaCAB* operon from *Ralstonia eutropha* H16. The hybrid *phaCAB* operon represents a step towards the more efficient production of P(3HB), which has an array of applications from 3D printing to tissue engineering.

## Introduction

Conventional oil-derived polyolefin plastics exhibit useful characteristics that have extensive commercial applications. However, the accumulation of plastics in the environment and the non-renewable source of polyolefin plastics have stimulated interest in sustainable sources of plastic production. Biopolymers, such as poly-3-hydroxy-butyrate (P(3HB)) are produced as a carbon store in an array of organisms and exhibit characteristics which are similar to oil-derived plastics^*(1, 2)*^. Furthermore, P(3HB) has the added advantages of biodegradability, biocompatibility and, based upon several life cycle analyses, P(3HB) production is more environmentally sustainable than polyolefin plastic production^*(1, 3, 4)*^. Despite these advantages, P(3HB) production is currently more expensive than the production of oil-derived plastics, and therefore more efficient P(3HB) production processes are required^*(5)*^. Genetic engineering approaches in which the P(3HB)-producing operon, *phaCAB*, is cloned into *Escherichia coli*, have pioneered the industrial production of P(3HB)^*(6)*^. More recently, synthetic biology approaches involving the rational engineering of the *phaCAB* operon^*(7)*^, and metabolic engineering strategies^*(8)*^ have continued to increase the efficiency of P(3HB) production. ‘*Team Plasticity’*, comprising undergraduate students from Imperial College London, set out to engineer a more efficient *phaCAB* operon as part of a project in the 2013 International Genetically Engineered Machine Competition (iGEM). The broader project aim was to exploit non-recyclable waste as a carbon source that could be utilized by engineered *E. coli* tasked with the *de novo* synthesis of P(3HB). In this paper we report on the characterization of a novel hybrid promoter *phaCAB* operon, which was generated during the project.

## Results and Discussion

The *phaCAB* operon from *Ralstonia eutropha* H16, is the most extensively studied P(3HB) synthesis operon^*(2)*^. It consists of three enzymes, which through a multi-stage enzymatic process generate P(3HB) inside the cell from the central metabolite acetyl-CoA^*(1)*^. The process is shown in Figure 1A and is briefly summarised here. Firstly, PhaA (3-ketothiolase) combines two molecules of acetyl-CoA to form acetoacetyl-CoA. Next, PhaB (acetoacetyl-CoA reductase) reduces acetoacetyl-CoA to form (R)-3-hydroxybutyl-CoA, which is then polymerised by PhaC (PHA synthase) to form poly-3-hydroxy-butyrate P(3HB). The *phaCAB* operon from *R. eutropha* H16 was originally cloned into *E. coli* in the late 1980s^*(2)*^. More recently the Tokyo Tech 2012 iGEM team (http://2012.igem.org/Team:Tokyo_Tech) cloned, characterised and standardized the native *phaCAB* operon into a biobrick-compatible format for the synthetic biology community.

**Figure 1.**
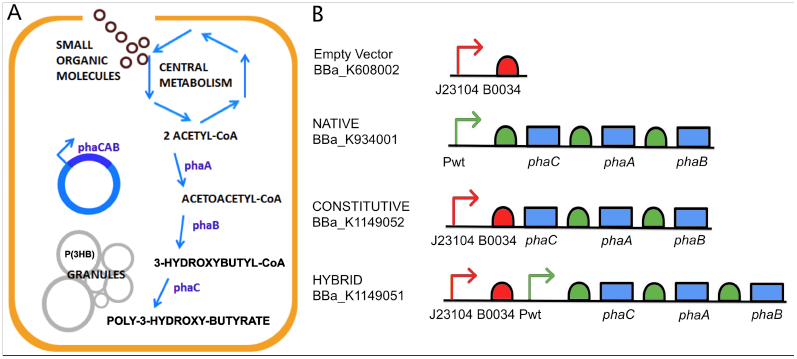
*phaCAB* pathway and constructs. (**A**) Schematic of poly-3-hydroxy-butyrate (P(3HB)) production via the *phaCAB* operon pathway. (**B**) The constructs utilized in this study. Abbreviations: Pwt (wildtype promoter; green arrow), J23104 (Anderson constitutive promoter, BBa_J23104; red arrow), B0034 (ribosomal binding site, BBa_B0034; red half-circle), *phaC* (PHA synthase), *phaA* (3-ketothiolase) and *phaB* (acetoacetyl-CoA reductase). Green half-circles denote native ribosomal binding sites. Construct symbols are based on the Synthetic Biology Open Language Visual (SBOLv) v1.0.0 guidelines.

In order to increase P(3HB) production, several novel *phaCAB* operons were engineered during the course of our project (Figure 1B). The constitutive *phaCAB* operon (BBa_K1149052) was designed such that the native promoter and RBS were replaced with a strong Anderson promoter (J23104) and the RBS B0034. Design considerations were based upon modeling simulations of the PhaCAB pathway in engineered *E. coli*. A sensitivity analysis of the synthetic pathway revealed that increasing expression of *phaB* would increase P(3HB) production. In order to increase expression of *phaB*, further simulations predicted that of the several designs that were tested, it was the constitutive operon design that would result in an increase in P(3HB) production (Figure 2). The hybrid operon design (BBa_K1149051) was constructed in parallel to the constitutive operon and is noted for its dual promoter and RBS combinations (Figure 1B).

**Figure 2.**
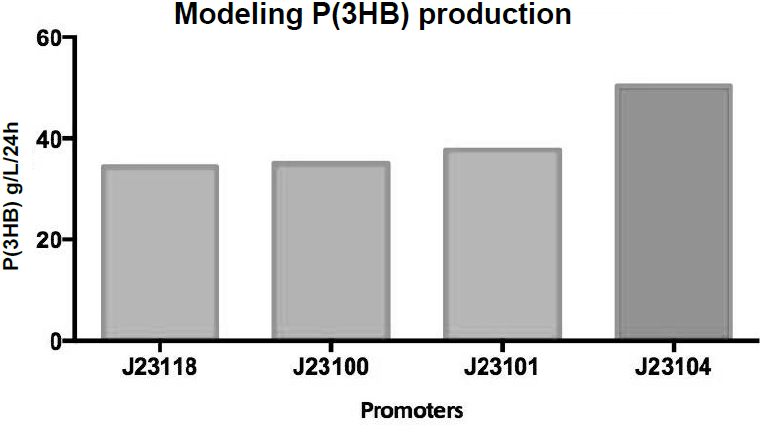
Modeling P(3HB) production. Simulation data showing P(3HB) production from different operon designs, where *phaCAB* expression is under the control of the indicated Anderson constitutive promoters.

*E. coli* MG1655 carrying either the empty vector (BBa_K608002), native *phaCAB* (BBa_K934001), constitutive *phaCAB* (BBa_K1149052) or hybrid *phaCAB* (BBa_K1149051) were cultured as described in Methods. P(3HB) was purified using sodium hypochlorite. Purified P(3HB) from each of the engineered populations were weighed and compared (Figure 3). Average P(3HB) production was around two-fold higher in constitutive *phaCAB*-engineered *E. coli* (0.49 g/L S.D. ± 0.06) compared to native *phaCAB* (0.22 g/L S.D. ± 0.18), while average P(3HB) production was six-fold higher in hybrid *phaCAB*-engineered *E. coli* (1.47 g/L S.D. ± 0.48), and empty vector-engineered *E. coli* did not produce detectable levels of P(3HB). Fluorescent microscopy and Nile Red staining further confirmed increased P(3HB) production. Full details are available on the iGEM Wiki, (http://2013.igem.org/Team:Imperial_College).

**Figure 3.**
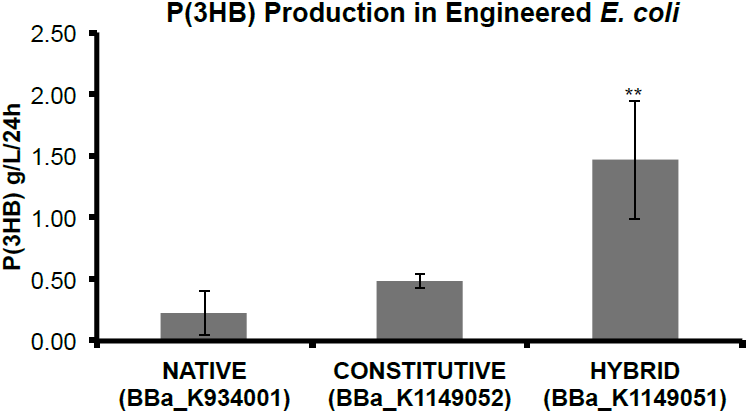
P(3HB) production in *E. coli* harboring *phaCAB* operon constructs. *E. coli* MG1655 transformed with either native (BBa_K934001), constitutive (BBa_K1149052) or hybrid (BBa_K1149051) *phaCAB* constructs were cultured in 1 liter LB media with 3% glucose (w/v) for 24 hours. P(3HB) was purified and weighed as described in Methods. Data represent the mean +/-the SD of three independent experiments. ** Student t-test, P <.01

It is likely that the dual promoter and RBS design (Figure 1B) of the hybrid system results in a higher level of mRNA transcript production and/or ribosome recruitment and thus an increase in the expression of the PhaCAB enzymes. Yet interestingly, P(3HB) production from the hybrid operon was greater than the combined P(3HB) production of the native and constitutive operons. Together, these data suggest that the hybrid promoter performs a multiplicative, rather than an additive combination of the native and constitutive promoter designs. Unlocking the design rules of the hybrid promoter may have applicability that extends beyond the *phaCAB* operon. Additionally, model-guided optimisation of the PhaCAB pathway led to the generation of the constitutive operon design, which in combination with the hybrid promoter, represent an emerging family of novel, rationally engineered P(3HB)-producing operons. These engineered operons in combination with additional synthetic biology approaches^*(7, 8)*^ and the switch towards low-cost carbon sources, such as mixed, non-recyclable waste, will further increase the efficiency and commercial viability of P(3HB) production.

## Methods

### Construct Assembly

Empty vector (BBa_K608002), and native *phaCAB* (BBa_K934001) constructs were sourced from the 2013 distribution of the iGEM Registry of Standard Biological Parts (partsregistry.org). Constitutive *phaCAB* (BBa_K1149052) was generated via PCR, with the native *phaCAB* operon (BBa_K934001) as the template. Primers Pha_Fw 5’-cgcttctagagatggctactgggaaaggagccg-3’ and BBa_G1005 5’-gtttcttcctgcagcggccgctactagta-3’, were utilised to generate a PCR product containing the *phaCAB* operon but excluding the native promoter and RBS. The PCR product was cloned into the destination vector (BBa_K608002) to generate the final constitutive *phaCAB* operon construct. The hybrid *phaCAB* (BBa_K1149051) construct was generated in parallel to the constitutive *phaCAB* operon. The entire native *phaCAB* operon including the native promoter and RBS was cloned into the destination vector (BBa_K608002) to generate the final hybrid *phaCAB* operon construct. All constructs were generated using the iGEM submission backbone, pSB1C3 and all restriction digests utilized the standard iGEM prefix and suffix restriction sites.

### P(3HB) production

*E. coli* strain MG1655 carrying either empty vector (BBa_K608002), native *phaCAB* (BBa_K934001), constitutive *phaCAB* (BBa_K1149052) or hybrid *phaCAB* (BBa_K1149051) constructs were grown in Luria Broth (LB) media supplemented with 34 µg/mL Chloramphenicol (final concentration) for maintenance of plasmids. Cultures were grown overnight at 37°C and 200 rpm shaking. Overnight cultures were diluted (1:200) into flasks containing 1 liter of LB, supplemented with 3% glucose (w/v) and 34 µg/mL Chloramphenicol. Cultures were subsequently grown for 24 h at 37°C and 200 rpm shaking, during which P(3HB) could be produced by *phaCAB* expressing E*. coli*.

### P(3HB) purification

1 liter production cultures were centrifuged at 4000 rpm for 15 minutes. Bacterial pellets were washed with phosphate-buffered saline (PBS) and then incubated with 1% Triton-X 100 in PBS (v/v) for 30 minutes at room temperature in order to lyse the cells. Final purification of P(3HB) utilized the sodium hypochlorite method as described in^*(9)*^.

### Modelling

The P(3HB) synthesis model was constructed and simulated using the Simbiology toolbox of Matlab. Full details are available on the iGEM Wiki (http://2013.igem.org/Team:Imperial_College)

## Acknowledgements

We would like to thank the members of the Centre for Synthetic Biology and Innovation (CSynBI), the companies Eppendorf, Bioline, GeneArt and additionally ERASysBio for sponsoring the 2013 Imperial College London iGEM project. We would also like to thank the iGEM Foundation and its volunteers for organizing the competition that instigated this work.

## Notes

The authors declare no conflict of interest and no competing financial interests.

